# Application of Microencapsulation Technology in Mushroom Powder Cake Processing

**DOI:** 10.1101/2020.11.24.395673

**Authors:** Liu Zifei, Ye Hongyan, Yue Wei, Ning Li

**Author notes:** College of Science and Engineering,Agricultural University of Hebei, Huanghua, China (Yue Wei). College of Food Science and Technology, Agricultural University of Hebei, Baoding, China (Ning Li). Corresponding author.(WY). These authors contributed equally to this work (Liu Zifei, Ye Hongyan).

## Abstract

In this study, *Lentinus edodes* (or mushroom) powder and carrot juice were used as raw materials to prepare a new type of nutritious cakes. Microencapsulation was applied in embedding vitamin C to prevent the loss of this vitamin during baking at high temperatures. The most suitable addition amount of each materials was optimized via single-factor test and response surface analysis. The optimal microencapsulation of shiitake mushroom powder with carrot juice required 10 g of mushroom powder, 35 g of soft sugar, 50 mL of carrot juice and 160 g of egg white. The cake prepared herein was rich in nutrient and exuded a unique flavor. This cake was prepared via modern processing technologies and efficient detection techniques. Microencapsulation not only improves the nutritional value of traditional flour products but also expands the scope of research on food processing technologies.

**Practical applications:** *Lentinus edodes* is a nutritious food with low fat content and rich in polysaccharides and vitamins. Consumption of this mushroom improves immunity, lowers blood pressure, and prevents the development of various types of cancer. Ground *L. edodes* powder is also called mushroom powder, which has the same nutrient composition and efficacy as *L.entinus edodes*. Carrots produce a fresh aroma, and the large amounts of β -carotene in carrots improve the nutritional value of carrot-based food and also exert beneficial effects, such as preventing xerophthalmia, night blindness, and oral ulcer. A new cake made of carrot juice and mushroom powder can effectively improve the nutritional and economic values of carrot and mushroom powder. However, vitamin C easily loses its function at high temperatures. In this study, vitamin C was microencapsulated to reduce its outflow. The purpose of this study was to expand the applications of microencapsulation technology, provide a scientific basis for the production of nutritious functional foods, and offer new research ideas for the development of new cakes.

## Introduction

*Lentinus edodes* (mushroom) is a nutritious food with high protein content, low fat content and rich in amino acids, polysaccharides, and vitamins [1]. It can improve immunity, lower blood pressure and blood lipid, prevent the development of cancer, and delay aging [2, 3]. Mushroom powder is obtained by cleaning the mushroom and removing its impurities with flowing water, drying it with hot air at 50 ℃ until its water content is lower than 14%, and crushing it into particles >120 μm in size [4, 5]. Mushroom powder can be used to make cakes or various flour products, such as steamed bread, cake, bread, biscuits, or crisp noodles. *L. edodes* powder has the same nutrients and beneficial effects as *L. edodes*, but it is not widely consumed because of its poor taste [6]. Carrot products a fresh aroma, and it can improve the taste of mushroom powder. Moreover, carrot is rich in β-carotene, and thus it can improve the nutritional value of carrot-based food [7, 8]. Carrot also has multiple beneficial effects, such as preventing dry eyes, night blindness, and oral ulcer [9, 10]. In recent years, some people have used carrots as ingredients for making cakes and other foods. However, at high temperatures, vitamin C easily loses its effectiveness. Previous studies suggested that the microencapsulation technology can retain the content of vitamin C in flour products and minimize its loss. In the present study, the microencapsulation technology was used to preserve the vitamin C content of cake made with carrot juice and mushroom powder.

The microencapsulation technology is a popular technology [11], extensively used in the food processing industry. This technology can preserve the biological activity of some functional ingredients and does not affect the color, flavor, taste, shape, and other characteristics of raw materials [12], especially beverages [13]. However, in-depth research on the production of flour products that require high temperatures is lacking [14].

In this study, carrot juice and *L. edodes* powder were used as ingredients to prepare and bake a cake. Vitamin C was microencapsulated to prevent its loss effectively during baking at high temperatures and preserve the content of vitamin C of the cake. The content of vitamin C was determined via high performance liquid chromatography (HPLC). Carrot juice was added to improve the taste and nutritional value of this cake. This study aimed to expand the applications of the microencapsulation technology, provide a scientific basis for the production of nutritious functional foods, and offer new research ideas for the development of new cakes.

## Materials and equipment

Mushroom powder, carrots, soft sugar, edible oil, eggs and baking powder are all purchased from a local supermarket. Vitamin C tablets were acquired from a local pharmacy. Food-grade sodium alginate and calcium lactate, L(+)-ascorbic acid standard (≥99% purity), D(-)-ascorbic acid standard (≥99% purity), methanol, and chromatographically pure hexadecyl trimethyl ammonium bromide were procured from Sigma-Aldrich Co. (St Louis, MO, USA). Other reagents used were analytically pure.

HPLC was performed using a 2998 ultraviolet detection HPLC system (Waters Corporation, of USA) with a lquid chromatographic column (5 um, 4.6mm × 250 mm;Atlantis^R^ T3). The raw materials were weighed using an electronic balance (Cixi Huaxu Weighing Instrument Industry Co., Ltd.). The raw materials were centrifuged using a high-speed refrigerated centrifuge (Anhui Zhongke Zhongjia Scientific Instrument Co., Ltd.). The egg beater, mixer, and oven were acquired from Guangdong Midea Household Appliances Manufacturing Co., Ltd.. Volume bottles and beakers were bought from Tianjin glass instrument Co., Ltd.

## Operating steps

### Process flow

Sodium alginate and vitamin C were evenly mixed to make a uniform solution. The solution was then dropped into a solidified calcium lactate solution to obtain the microencapsulated finished product. *L. edodes* powder, cake powder, carrot juice, edible oil, baking powder, soft sugar, and egg yolk were mixed to make a batter. Egg white were added, and the microencapsulated finished product was added into the batter. The batter was poured into a mold and left in an oven to bake. The process is depicted (Fig 1).

**Fig 1.** Process flow diagram.

### Pre-experimental operation steps

#### Preparation of microcapsules

First, 0.75 g of sodium alginate was added to 50 g of purified water. The mixture was stirred until clear and set aside for later use. Afterward, 0.2 g of vitamin C tablet was ground into powder and then added into the sodium alginate solution. Subsequently, 2 g of calcium lactate was added in 50 g of purified water. The mixture was stirred until clear, and set aside for later use. Finally,the mixed solution of sodium alginate and vitamin C was dripped into the solidified calcium lactate solution by using a dropper to form spherical microcapsules.

#### Preparation of carrot juice

Carrots were cleaned with water and then cut into pieces, several of which were put in a juicer. Twice the volume of purified water was poured into the juicer, and then the carrots were squeezed. The juice was set aside for later use.

#### Blending of batter

*L. edodes* powder and cake powder were mixed at a certain proportion. An appropriate volume/amount of carrot juice, soft sugar, edible oil, baking powder, and beaten egg yolks and egg white were added to the powder. The mixture was evenly mixed to obtain the batter.

#### Injection molding

The batter was poured into the cake crust, accounting for about 2/3 of the cake crust. An appropriate amount of microcapsules was added, and the batter was mixed well.

#### Baking

The oven was preheated at 180 ℃ for about 5 min. Afterward, the cake was placed in the oven at 180 ℃ and baked for 20 min to obtain the finished product.

### Experimental design

#### Single-factor tests

According to the results of our preliminary experiment, a single-factor experiment was conducted with the addition amount of mushroom powder, soft white sugar, carrot juice and egg white as the factors.

First, 30 g of soft sugar, 40 mL of carrot juice, 20 mL of edible oil, 40 g of egg yolk, 80 g of egg white, and 50 g of flour were mixed. The mixture was then baked at 180 ℃ for 20 min. Different amounts of mushroom powder (5, 10, 15, 20, and 25 g) were added. The rest was cake powder. Finally, the optimal addition amount of mushroom powder was determined according to physicochemical and sensory indexes. The above operations show how to optimize the addition amount of mushroom powder.

Different amount of soft sugar (25, 30, 35, 40, and 45 g) were added.Afterward, 10 g of mushroom powder, 40 g of cake powder, and the addition amount of other raw materials and baking operations are the same as those in the previous step. The optimal addition amount of soft sugar was determined according to physicochemical and sensory indexes. The above operations show how to optimize the addition amount of soft sugar.

Different amounts of carrot juice (20, 30, 40, 50, and 60 mL), 10 g of mushroom powder, 40 g of cake powder, and the addition amount of other raw materials and baking operations are the same as those in the previous step. The optimal addition amount of carrot juice was determined according to physicochemical and sensory indexes. The above operations show how to optimize the addition volume of carrot juice.

Different amounts of egg white (40, 80, 120, 160, and 200 g), 10 g of mushroom powder, 40 g of cake powder, and the addition amount of other raw materials and baking operations are the same as those in the previous step. The optimal addition amount of egg white was deteremined according to physicochemical and sensory indexes. The above operations show how to optimize the addition amount of egg white.

#### Response surface optimization of experimental conditions

On the basis of the results of single-factor tests results, sensory scores and vitamin C contents were taken as the response values. The effects of the main influencing factors on the sensory scores and vitamin C contents were evaluated using the Design-Expert 8.0.6 software. The microencapsulation processing technology was optimized. The test factor levels [15, 16]are summarized (Table 1).

**Table 1.**
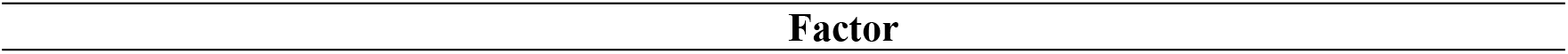
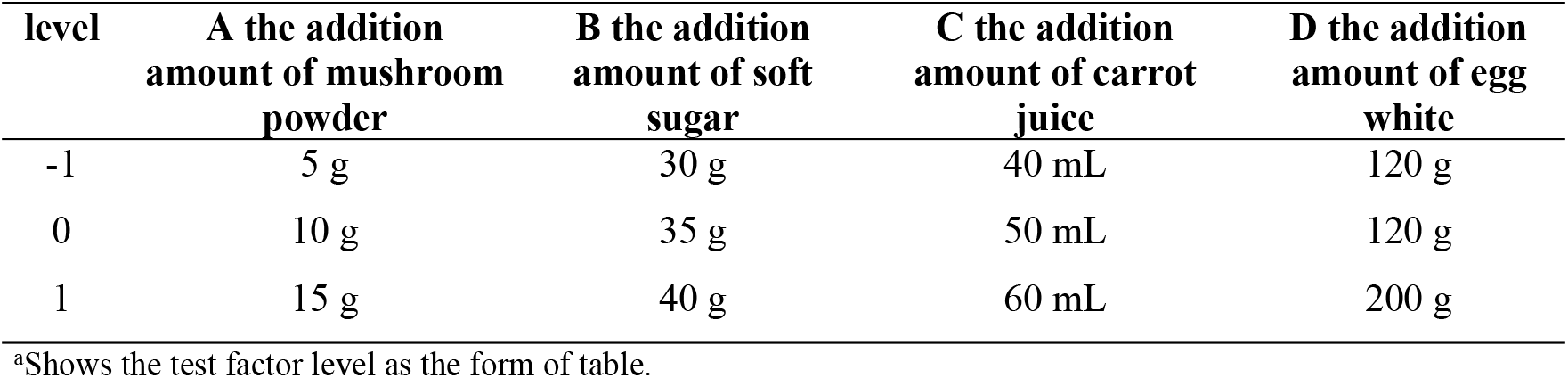
Level table of response surface analysis test factors.

| level | A the addition<br>amount of mushroom<br>powder | B the addition<br>amount of soft<br>sugar | C the addition<br>amount of carrot<br>juice | D the addition<br>amount of egg<br>white |
| --- | --- | --- | --- | --- |
| -1 | 5 g | 30 g | 40 mL | 120 g |
| 0 | 10 g | 35 g | 50 mL | 120 g |
| 1 | 15 g | 40 g | 60 mL | 200 g |

<sup>a</sup>Shows the test factor level as the form of table.

#### Product quality evaluation

Ten professionals were selected to evaluate the cake on five sensory indexes of color, appearance, taste, organization, and smell.Their scores were averaged. The criteria are listed (Table 2).

**Table 2.** Grading standard of mushroom powder microencapsulated cake.

| Projects | Scoring rules | Score |
| --- | --- | --- |
| Color | surface color: golden yellow | 11~15 |
|  | surface color : yellow | 6~10 |
|  | the surface has irregular caramel color | 0~5 |
| Appearance status | full, normal uplift | 15~20 |
|  | There is a slight deformation. | 8~14 |
|  | not full and cracked | 0~7 |
| Taste | soft and soft in taste | 21~30 |
|  | The taste is not soft and sticky. | 11~20 |
|  | The taste is not soft and has obvious rough feeling. | 0~10 |
| Organization | fine pores and no cracks | 11~15 |
|  | stomata slightly thick, some cracks | 6~10 |
|  | pores are coarse and cracked. | 0~5 |
| Smell | has a unique cake smell | 15~20 |
|  | odorlessness | 8~14 |
|  | odor | 0~7 |

Physical and chemical indexes include the content of vitamin C in cake and the specific volume of cake.

The vitamin C content of the cake was determined according to GB 5009.86-2016 guidelines [17]. Ultrapure water as mobile phase A and methanol as mobiles phase B were mixed at a ratio of 98: 2. The mixture was filtered through a 0.45 μm water phase filter membrane. The vitamin C content of the finished product was determined after ultrasonic degassing.

Specific volume was measured using formula 1:

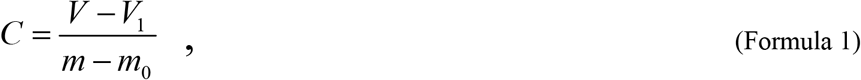

where *m_0_* is the mass of the mold, *m* is the mass of the finished product after cooling for 30 min, *V_1_* is the original volume of millet, and *V* is the volume of cake after adding millet. (Volume was measured via the millet method [18].)Each sample was measured in parallel three times, and the specific volume score of the cake was taken as the average value.

The specific volume scores of the cake according to GB/T 24303-2009 “Inspection of grain and oils: Method for cake-making of wheat flour-Sponge cake” are summarized (Table 3).

**Table 3.** Cake specific volume rating table.

| Specific volume<br>/(mL/g) | Score | Specific volume<br>/(mL/g) | Score | Specific volume<br>/(mL/g) | Score | Specific volume<br>/(mL/g) | Score |
| --- | --- | --- | --- | --- | --- | --- | --- |
| 2.5 | 7 | 3.4 | 16 | 4.3 | 25 | 5.2 | 26 |
| 2.6 | 8 | 3.5 | 17 | 4.4 | 26 | 5.3 | 25 |
| 2.7 | 9 | 3.6 | 18 | 4.5 | 27 | 5.4 | 24 |
| 2.8 | 10 | 3.7 | 19 | 4.6 | 28 | 5.5 | 23 |
| 2.9 | 11 | 3.8 | 20 | 4.7 | 29 | 5.6 | 22 |
| 3.0 | 12 | 3.9 | 21 | 4.8 | 30 | 5.7 | 21 |
| 3.1 | 13 | 4.0 | 22 | 4.9 | 29 | 5.8 | 20 |
| 3.2 | 14 | 4.1 | 23 | 5.0 | 28 | 5.9 | 19 |
| 3.3 | 15 | 4.2 | 24 | 5.1 | 27 | 6.0 | 18 |

<sup>a</sup>Shows the scoring standard of cake specific volume in GB/T 24303-2009.

## Results and analysis

### Single factor experiment results

#### Optimization of *L. edodes* powder addition amount

The physicochemical and sensory indexes of different addition amounts of *L. edodes* powder(5, 10, 15, 20, and 25 g) were measured to determine the optimal addition amount of the mushroom powder to add(Fig. 2).

**Fig 2.** Effect of the addition amount of mushroom powder on cake quality.

The specific volume scores and vitamin C contents initially increased and then decreased as the amount of mushroom powder added increased (Fig. 2). The specific volume score was the highest when 10 g of the mushroom powder was added. At this moment, the cake had the best fluffiness and softest taste. The vitamin C content was the highest when 15 g of the mushroom powder was added. Therefore, microencapsulation effectively embedded the vitamin C of the cake. In spite of the high temperature at which the cake was baked, its vitamin C content was preserved to the greatest extent, indicating that vitamin C was protected and a large loss of vitamin C content was prevented. The sensory scores initially increased and then decreased as the addition amount of *L. edodes* powder increased. The sensory score was the highest when the addition amount of the mushroom powder was 10 g, but it gradually decreased.*L. edodes* powder itself has a unique flavor, but it does not taste good. When the addition amount was less than 10 g, the unique flavor of *L. edodes* was not obvious. However, the cake tasted bad and the smell of the mushroom was strong when excessive amounts of the mushroom powder were added. Therefore, the optimal addition amount of *L. edodes* powder was 10 g.

#### Optimization of soft sugar addition amount

The physicochemical and sensory indexes of different addition amounts of soft sugar (25, 30, 35, 40, and 45 g) were valuated (Fig. 3).

**Fig 3.** Effect of the addition amount of soft sugar on cake quality.

The specific volume scores and vitamin C contents initially increased and then decreased as the addition amount of soft sugar increased (Fig.3). The specific volume scores and vitamin C contents were the highest when 35 g of soft sugar was added. However, microencapsulation had negligible effectes on the specific volume scores and vitamin C contents of the cake. Nevertheless, the cake had the best fluffiness. The microencapsulation technology was adopted to preserve the vitamin C content of the cake as much as possible.The cake was microencapsulated, to protect vitamin C effectively and prevent large losses of this vitamin at high temperatures. The sensory score was highest when 35 g of soft sugar was added, but it gradually decreased. Adding soft sugar increased the sweetness of the cake and improved its taste. When the addition amount of soft sugar was less than 35 g, the sweetness of the cake was low and it was tasteless. When the addition amount exceeded 35 g, the sweetness increased, but the cake felt greasy and it tasted bad. Therefore, the optimal addition amount of soft sugar to add was 35 g.

#### Optimization of carrot juice addition amount

The physicochemical and sensory indexes of different volumes of carrot juice added (20, 30, 40, 50, and 60 mL) were determined, (Fig. 4).

**Fig 4.** Effect of the addition volume of carrot juice on cake quality.

The specific volume scores initially increased and then decreased as the addition volume of carrot juice increased; by comparison, the vitamin C content of the cake increased. The specific volume score was the highest, and the cake had the best fluffiness and softest taste when the addition volume of carrot juice was 40 mL. The vitamin C content was the highest when the addition volume was 60 mL. Thus, microencapsulation effectively embedded vitamin C, and protected it at high temperatures, thereby preventing large losses of vitamin C during baking at high temperatures.The sensory score of the cake initially increased and then decreased as the addition volume of carrot juice increased. When the addition volume was 50 mL, the sensory score was the highest, but it gradually decreased. Adding carrot juice made the cake more fragrant and improved its taste. When an insufficient volume of carrot juice was added, the fragrant taste of carrot juice was not obvious and the carrot flavor could not be tasted. By contrast, when excessive volumes of carrot juice were added, the cake tasted heavy, and the people we asked to taste it did not like the cake. Therefore, the optimal addition volume of carrot juice to add was 50 mL.

#### Optimization of addition amounts of egg white

The physicochemical and sensory indexes of different amounts of egg white added (40, 80, 120, 160, and 200 g) were determined, (Fig. 5).

**Fig 5.** Effect of the addition amount of egg white on cake quality.

The specific volume scores increased as the addition amount of egg white increased. By comparison, the vitamin C content initially increased and then decreased. The specific volume score was the highest when the addition amount of egg white was 200 g. By comparison, the vitamin C content was the hightest when the addition amount of egg white was 120 g. Thus, microencapsulation effectively embedded vitamin C and preseved the vitamin C content of the cake as much as possible despite the high temperatures. Microencapsulation effectively protected vitamin C, ensuring that the cake would retain this vitamin. The sensory score was the highest when 160 g of egg white was added, but it gradually decreased.The addition of egg white made the cake fluffier, softer, and with a better taste. When the addition amount of egg white less than 160 g, the finished cake was not fluffy. When too much egg white were added, the cake was too fluffy and tasted bad. Therefore, the optimal addition amount of egg white to add was 160 g.

### Analysis of the response surface optimization tests

#### Results of response surface tests

Sensory scores and vitamin C contents were taken as the response values in single-factor experiments.The response values were analyzed, using the Box–Behnken model in Design-Expert 8.0.6 software (Table 4).

**Table 4.** Experimental results of response surface analysis.

| Experimental number | Addition amount of mushroom powder A | Addition amount of soft sugar B | Addition amount of carrot juice C | Addition amount of egg white added D | Sensory score (points) Y<sub>1</sub> | VitaminC content (mg/100g) Y<sub>2</sub> |
| --- | --- | --- | --- | --- | --- | --- |
| 1 | 0 | 0 | 0 | 0 | 93.6 | 93.01 |
| 2 | 0 | 0 | 0 | 0 | 94 | 92.98 |
| 3 | – 1 | 0 | 0 | 1 | 86.6 | 88.25 |
| 4 | 0 | 0 | 0 | 0 | 93.5 | 91.97 |
| 5 | 0 | 1 | 1 | 0 | 89.5 | 90.52 |
| 6 | 1 | 1 | 0 | 0 | 90.5 | 91.94 |
| 7 | 0 | 0 | 1 | – 1 | 87 | 88.87 |
| 8 | – 1 | 1 | 0 | 0 | 88 | 89.23 |
| 9 | 0 | 0 | - 1 | 1 | 84.7 | 86.21 |
| 10 | 1 | 0 | 1 | 0 | 89.5 | 90.87 |
| 11 | - 1 | 0 | 0 | - 1 | 88.5 | 89.12 |
| 12 | 0 | 0 | - 1 | - 1 | 86.5 | 87.82 |
| 13 | 1 | 0 | 0 | 1 | 89.5 | 90.85 |
| 14 | 0 | 1 | 0 | 1 | 85.4 | 85.5 |
| 15 | 0 | 0 | 0 | 0 | 93.2 | 92.45 |
| 16 | 0 | - 1 | 0 | - 1 | 85.6 | 85.98 |
| 17 | 0 | - 1 | 1 | 0 | 87.5 | 88.96 |
| 18 | 1 | - 1 | 0 | 0 | 87.5 | 89.09 |
| 19 | 0 | 1 | 0 | - 1 | 90 | 91.39 |
| 20 | 0 | 0 | 0 | 0 | 93.5 | 92.19 |
| 21 | 0 | - 1 | - 1 | 0 | 86 | 88.15 |
| 22 | 1 | 0 | - 1 | 0 | 89 | 89.712 |
| 23 | 0 | - 1 | 0 | 1 | 88 | 89.53 |
| 24 | 0 | 0 | 1 | 1 | 89 | 90.09 |
| 25 | - 1 | 0 | - 1 | 0 | 86 | 87.52 |
| 26 | 1 | 0 | 0 | - 1 | 88.9 | 89.79 |
| 27 | - 1 | - 1 | 0 | 0 | 89.5 | 90.29 |
| 28 | - 1 | 0 | 1 | 0 | 90 | 91.27 |
| 29 | 0 | 1 | - 1 | 0 | 87.5 | 87.89 |

<sup>a</sup>Shows the analysis results of the single factor.

#### Analysis of variance

According to the results of response surface tests and with sensory evaluation as the response value, the regression equation was determined as follows [19, 20]:

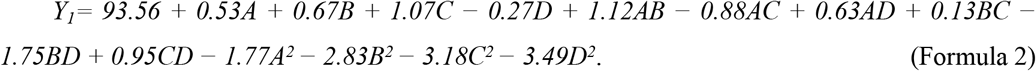

The order of importance of the main influencing factors was as follows: the volume of carrot juice > the amount of soft sugar > the amount of mushroom powder > the amount of egg white (Table 5). The F-value of model Y_1_ was 63.49 (p < 0.0001), indicating that differences in these factors were very significant. Misfitting term p was 0.1337 > 0.05, and no significant differences were observed in the misfitting term. The correlation coefficient of model Y_1_ was 0.9845, which explained 98.45% of the charge in response value.The equation was tested and results showed that the primary term (C), the interaction term of soft sugar addition amount and egg white addition amount (BD), the quadratic term of mushroom powder addition amount (A^2^), the quadratic term of soft sugar addition amount (B^2^), the quadratic term of carrot juice addition amount (C^2^), and the quadratic term of egg white addition amount (D^2^) significantly affected the sensory scores (p < 0.0001). Moreover, amounts of *L. edodes* powder (A),amounts of soft sugar (B), interaction items of *L. edodes* powder and soft sugar (AB), interaction items of *L. edodes* powder and carrot juice (AC), interaction items of carrot juice and egg white (CD) significantly affected the sensory scores (p < 0.01). Furthermore, the interaction item of mushroom powder and egg white (AD) significantly affected the sensory scores (p < 0.05). By contrast, the other variables had no significant effects on the sensory scores (p > 0.05).

**Table 5.**
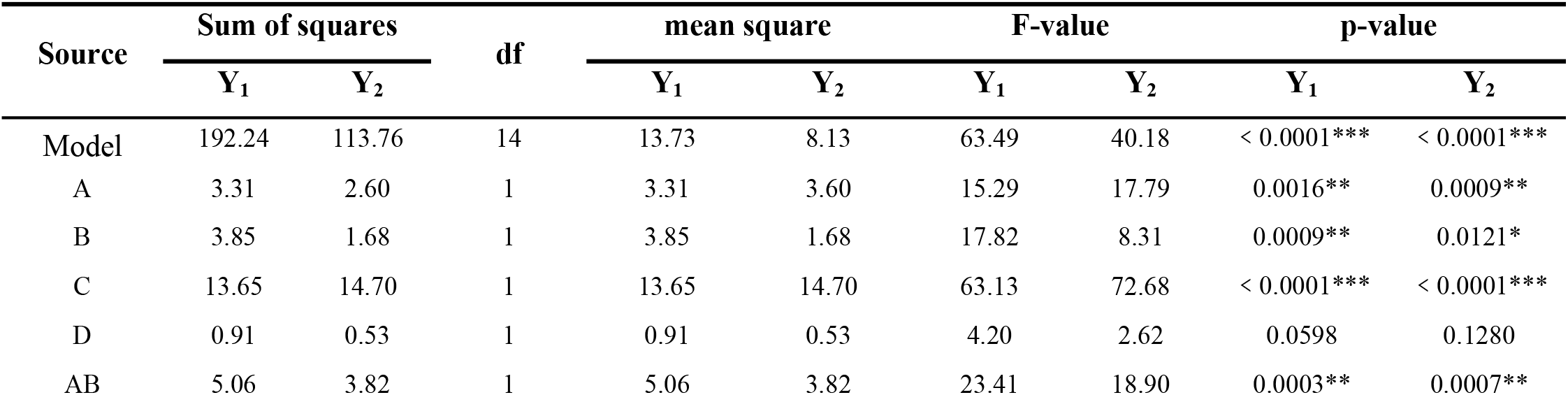
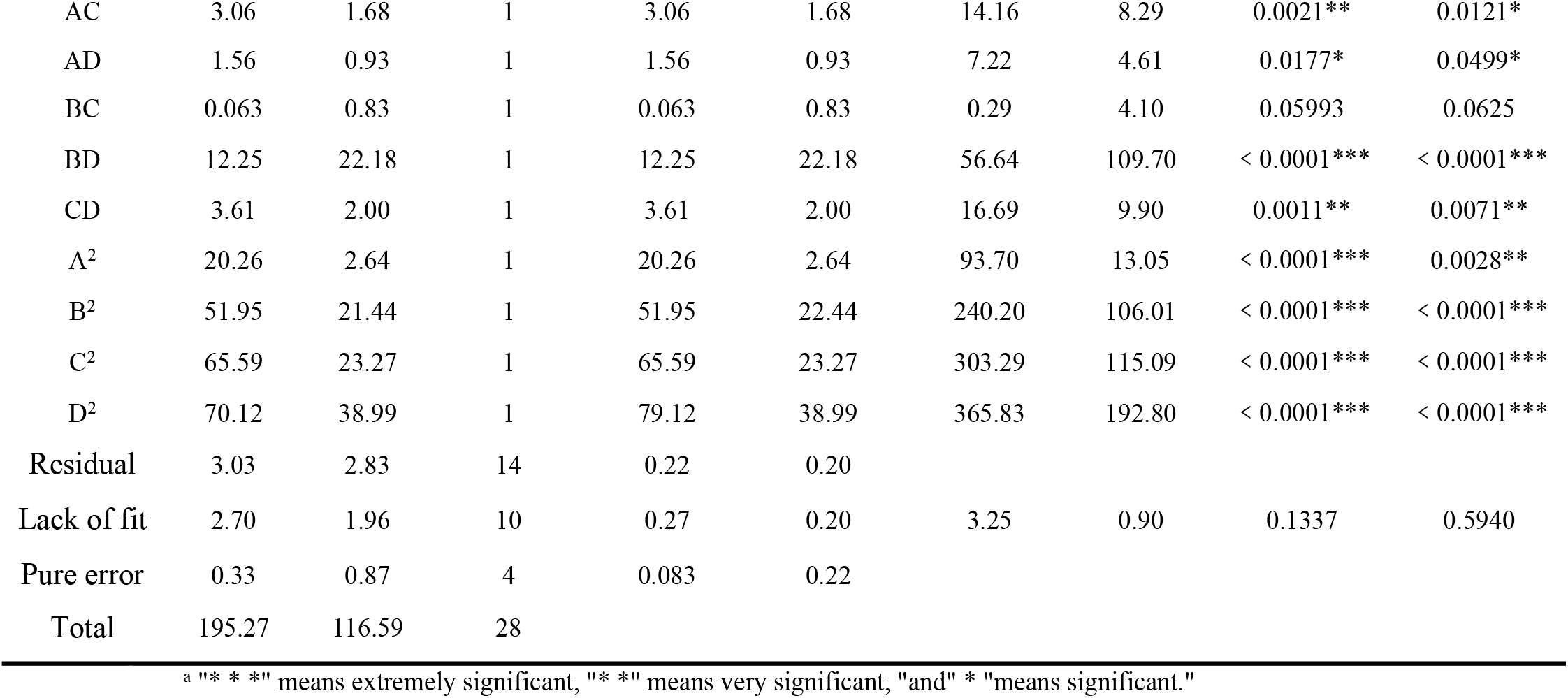
Variance analysis of regression equation.

With vitamin C content as the response value, the regression equation was determined as follows:

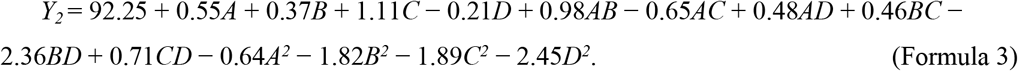

The order of importance of the influencing factors was as follows: the volume of carrot juice > the amount of mushroom powder > the amount of soft sugar > the amount of egg white (Table 5). The F-value of Y_2_ was 40.21 (p < 0.0001), indicating that model Y_2_ was extremely significant. Misfitting items p were 0.5943 > 0.05, and no significant differences were observed in the misffiting items. The correlation coefficient of model Y_2_ was 0.9757, indicating that this model explained 97.57% of the change in response value. The significance test of the regression equation showed that the primary term (C), the interactive term (BD), and the quadratic terms (B^2^, C^2^, and D^2^) of soft sugar in the model significantly affected the vitamin C contents (p < 0.0001). The primary term A, the interactive terms AB and CD, and the quadratic term A^2^ significantly influenced the vitamin C contents (p < 0.01). The primary item B and the interactive items AC and AD significantly influenced the vitamin C contents (p < 0.05). The other variables had no significant effects on vitamin C contents (p > 0.05).

#### Transactional analysis among factors

With sensory scores and vitamin C contents as the response values, the interaction response surface of each factor was designed using the Design-Expert 8.0.6 software. Response surface diagram and contour line plots [21] are illustrated in Figs. 6 and 7, respectively.

**Fig 6.** Effects of various factors on sensory scores of cake. This set of figures include response surface diagrams and contour line plots. (a) shows the interactive effect of mushroom powder addition amount and soft sugar addition amount on sensory scores of cake. (b) shows the interactive effect of mushroom powder addition amount and carrot juice addition volume on sensory scores of cake. (c) shows the interactive effect of mushroom powder addition amount and egg white addition amount on sensory scores of cake. (d) shows the interactive effect of carrot juice addition volume and soft sugar addition amount on sensory scores of cake. (e) shows the interactive effect of soft sugar addition amount and egg white addition amount on sensory scores of cake. (f) shows the interactive effect of carrot juice addition volume and egg white addition amount on sensory scores of cake.

**Figure 7.** Effects of various factors on vitamin C content of cake. This set of figures include response surface diagrams and contour line plots, (a) shows the interactive effect of mushroom powder addition amount and soft sugar addition amount on vitamin C content in cake. (b) shows the interactive effect of mushroom powder addition amount and carrot juice addition volume on vitamin C in cake. (c) shows the interactive effect of mushroom powder addition amount and egg white addition amount on vitamin C content in cake. (d) shows the interactive effect of carrots juice addition volume and soft sugar addition amount on vitamin C content in cake. (e) shows the interactive effect of soft sugar addition amount and egg white addition amount on vitamin C content in cake. (f) shows the interactive effect of carrot juice addition volume and egg white addition amount on vitamin C content in cake.

Response surface diagram can directly reflect the influence of the interactions of various factors on response values. The steeper the response curves are, the more elliptical the contour graphs will be, indicating strong interactions. The contour line plots of a, b, c, e, and f in Fig. 6 were oval, and the curve line plot in Fig. 6e had the largest slope. The interaction between soft sugar addition amount and egg white addition amount (BD) had a substantial effect on the sensory scores, whereas the interaction of addition amount soft sugar and addition volume of carrot juice (BC) had no considerable effect on the sensory scores. According to the p values, the order of importance of the interactive factors affecting the sensory scores was BD > AB > CD > AC > AD > BC. When the other factors were at the 0 level, the sensory scores initially increased and then decreased as the addition amount of mushroom powder, soft sugar, and egg white and the addition volume of carrot juice increased. The maximum sensory scores was reached when each factor was at the 0 level. At this point, the cake was golden yellow, had a fine taste, had fine pores, and exuded the unique flavors of carrot and mushroom. Therefore, there were extreme values in the selected parameters, which can predict the best technology of cake.

The contour line plot was round in Fig. 7d, indicating that the interaction between the addition amount of soft sugar and the addition volume of carrot juice (BC) had no significant influence on vitamin C content. By contrast, the interaction of other factors had a significant influence on the vitamin C content in cake. According to the p values, the order of importance of the influence of each interactive factor on vitamin C content in cake was BD > AB > CD > AC > AD > BC. When the other factors were at the 0 level, the vitamin C content of the cake initially increased and then decreased as the addition amount of mushroom powder, soft sugar, and egg white and the addition volume of carrot juice increased. The maximum vitamin C content was reached when all factors were at the 0 level, indicating that the vitamin C content of the cake was the highest, which demonstrated that microencapsulation preserved the vitamin C content of the cake to the greatest extent, ensuring the nutritional value of the cake. The present results [22] were consistent with those of Lee et al., who reported that microencapsulation can effectively prevent the loss of vitamin C even at high temperatures. The results demonstrated the feasibility and effectiveness of embedding vitamin C by microencapsulation at high temperatures. Microencapsulation ensures the nutritional value of food and broadens potential applications of this technology.

### Verification Tests

With sensory scores and vitamin C contents as the response values, response surface analysis revealed that the optimal addition amounts of the ingredients were 11.19 g of *L. edodes* powder, 35.95 g of soft sugar, 51.79 mL of carrot juice, and 158.56 g of egg white. Under optimal conditions, the predicted sensory scores were 93.7019, and the vitamin C content was 92.8057 mg/100 g. Actual baking conditions required that the addition amounts of mushroom powder, soft sugar, and egg white and the addition volume of carrot juice to add were 10, 35, and 160 g and 50 mL, respectively. The actual sensory scores were 93, and vitamin C content was 92.451 mg/100 g. The deviation between the predicted and actual values was small, confirming the effectiveness of the models.

Zhang et al. [23] explored the best processing conditions for baking carrot filaments. The vitamin C content of the cake was 12 mg/100 g. Unlike this study, the present work adopted the microencapsulation technology to embed vitamin C and used carrot and mushroom powder as raw materials to prepare and bake the cake. The vitamin C content of the final cake was 92.451 mg/100 g. This content was evidently higher than that of pure carrot filament cake without vitamin C embedding. This study demonstrated that the vitamin C content of cakes can be effectively improved by using the microencapsulation technology in embedding vitamin C. The results verified the feasibility of embedding vitamin C in baking cakes by using the microencapsulation technology. This study broadens the potential applications of the microencapsulation technology.

## Product quality evaluation

### Sensory index

The cake surface is golden yellow in color, normal in appearance and plump, with delicate taste, fine and uniform pore structure, no cracks on the surface, and unique smell of carrot and mushroom. The pores inside the cake were fine. Results of sensory evaluation showed that the addition volume of carrot juice positively affected the sensory indexes of the cake. Moreover, the addition volume of carrot juice improved the nutritional value of the cake, complemented the unique smell of the mushroom powder, provided fresh aroma, and enhanced the taste of the cake. These positive features make this type of cake palatable to consumers.

### Physical and chemical indicators

The specific volume scores were 27 points, and the vitamin C content of the cake was 44.6875 mg/100 g.

The specific volume scores of the cake were extraodinary ideal, and the cake was fluffy. Unlike the traditional way of baking carrot and mushroom powder cake, the cake prepared and baked herein used carrot and mushroom powder as raw materials, and vitamin C was embedded via the microencapsulation technology to prevent the loss of vitamin C during baking at high temperatures. Microencapsulation preserved the vitamin C content of the cake despite high temperatures. Thus, this technology improved the economic and nutritional value of the cake. This study broadens the potential applications of microcapsulation technology for use in flour products.

## Conclusion

In this study, vitamin C was embedded via the microencapsulation technology. Microencapsulation effectively prevented the loss of vitamin C during baking at high temperatures. Moreover, this new type of cake was made with mushroom powder and carrot as the main raw materials. These materials also enhanced the content of vitamin C of the cake. The prevention of the loss of vitamin C at high temperatures during baking improved the nutritional value and economic value of the cake. Single-factor tests and response surface analysis revealed that the order of importance of the main influencing factors was volume of carrot juice > amount of soft sugar > amount of mushroom powder > amount of egg white. The best processing conditions in actual baking required 10 g of mushroom powder, 35 g of soft sugar, 50 mL of carrot juice, and 160 g of egg white. The vitamin C content of the cake was determined. Results showed that the vitamin C content of the cake prepared via the microencapsulation technology was substantially higher than that of pure carrot cake without microencapsulation. The cake prepared and baked herein tasted good, exuded a unique flavor, and had a high nutritional and economic value. This cake broadens the market for mushroom powder- and carrot-based food. Moreover, this study expands the potential applications of the microencapsulation technology for use in flour products and offers new research ideas for the development of new types of cake.

## Acknowledgements

This project is supported by the Hebei Agricultural University Science and Technology Fund (C2020204129). In the process of research and topic selection, this experiment has been guided by President Ma and Teacher Wei. Thanks to teachers, every data and experimental details of this article can not be separated from the teachers’ teaching. Thanks to the school for providing laboratory and experimental equipment for the search. Thanks to Hebei Agricultural University for training and educating us.

## Supporting information

**S1 Fig. 1.**
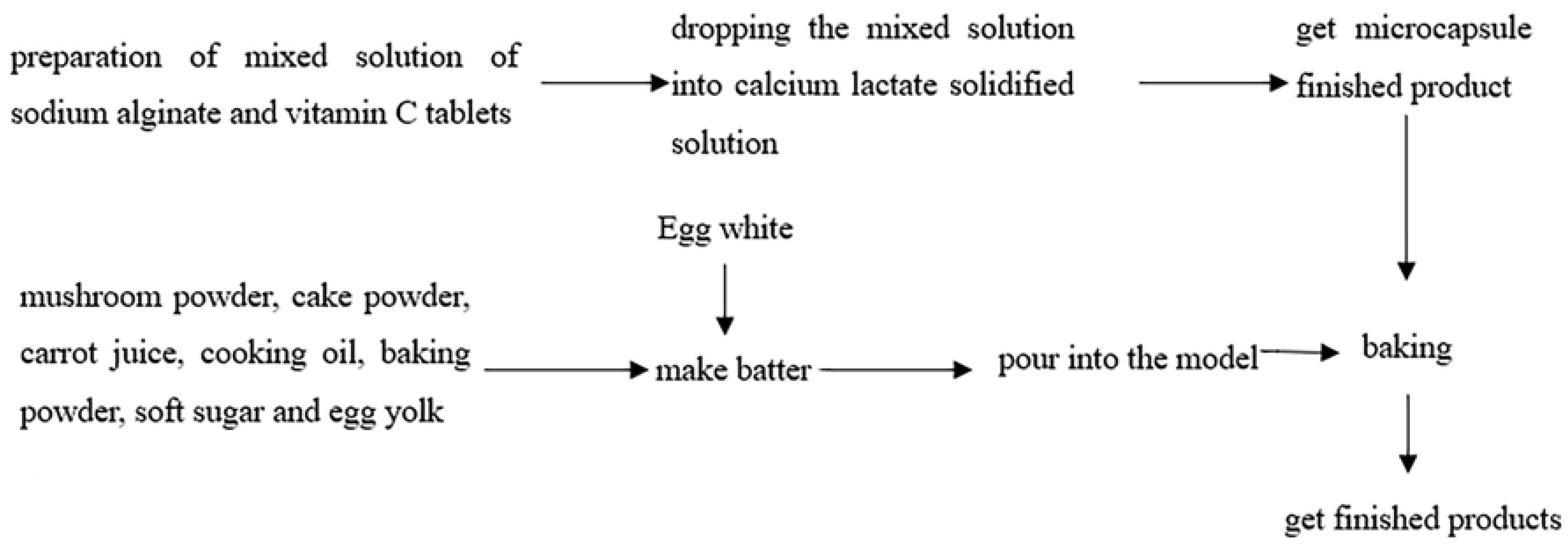
Process flow diagram.

**S1 Table. Level table of response surface analysis test factors.** The table shows the test factors level as the form of table.

**S2 Table. Grading standard of mushroom powder microencapsulated cake.**

**S3 Table. Cake specific volume rating table.** The table shows the scoring standard of cake specific volume in GB/T 24303-2009.

**S2 Fig. 2.**
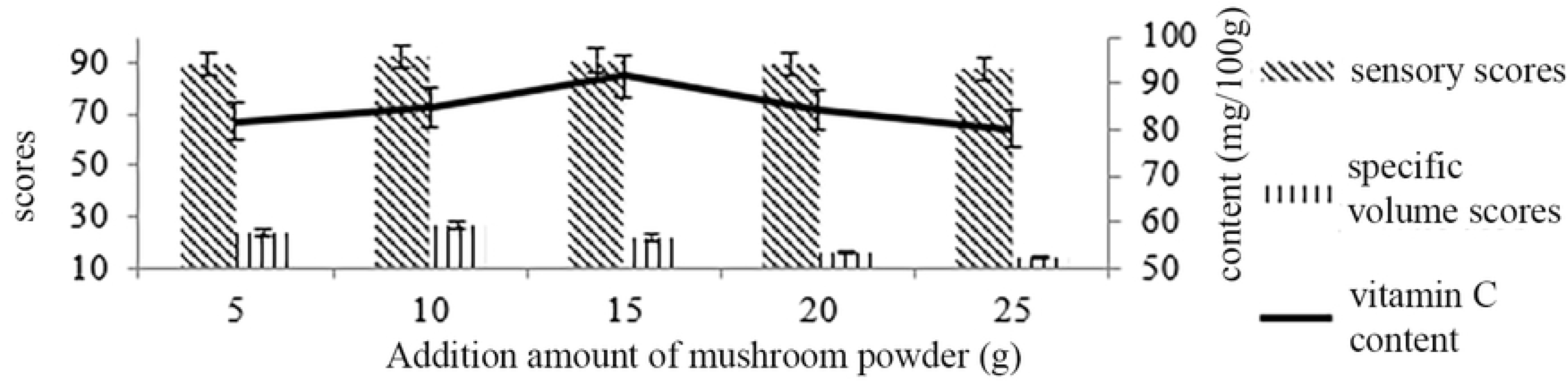
Effect of mushroom powder on cake quality.

**S3 Fig. 3.**
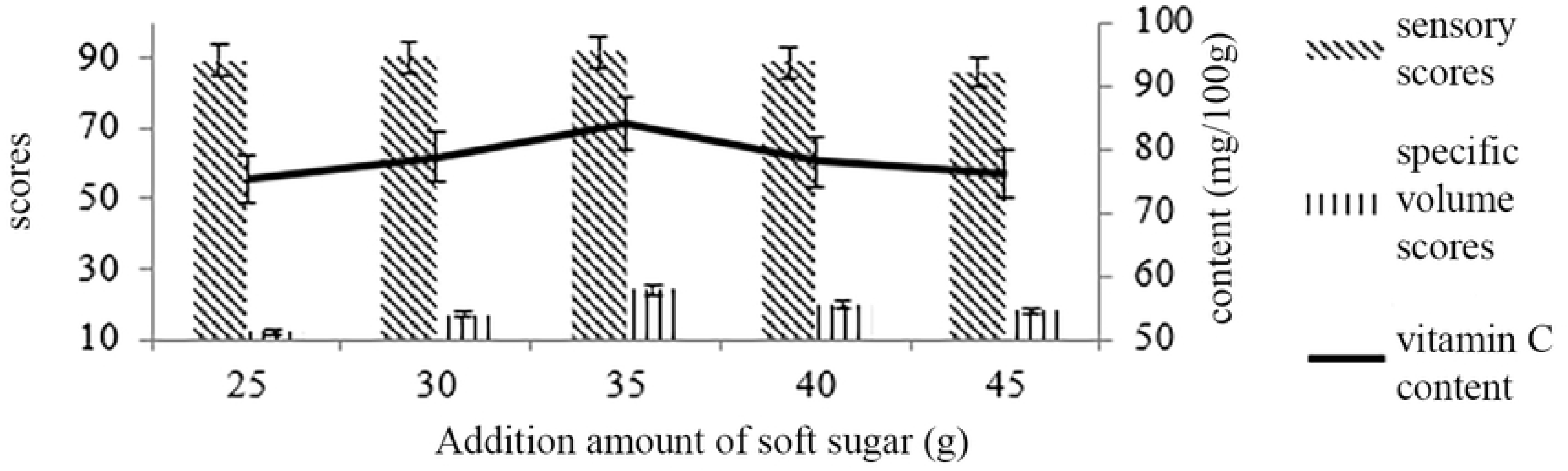
Effect of the amount of soft sugar on cake quality.

**S4 Fig. 4.**
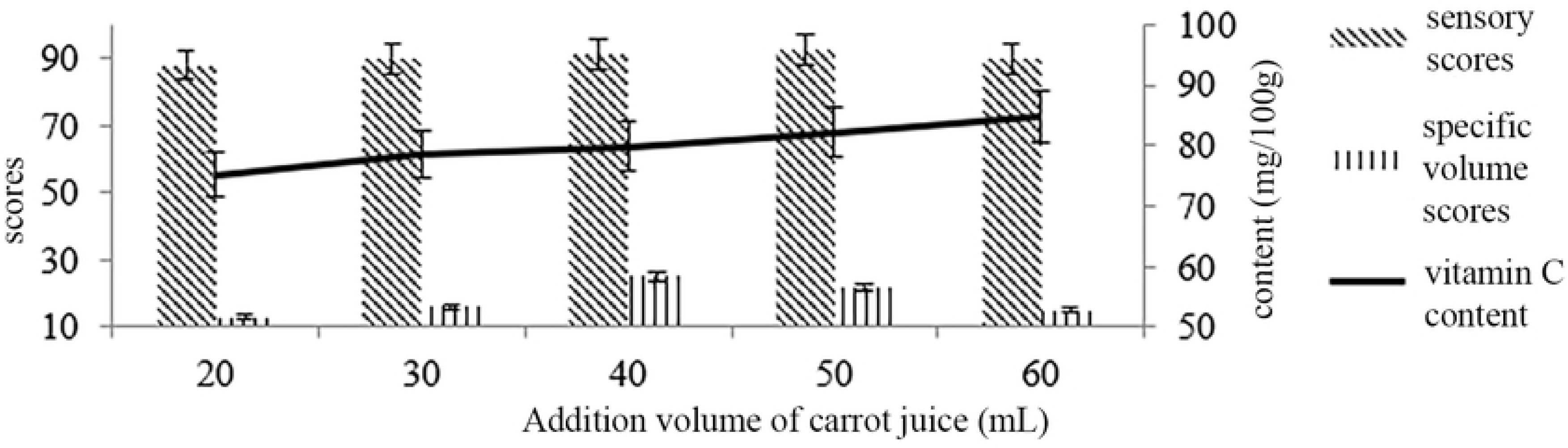
Effect of the volume of carrot juice on cake quality.

**S5 Fig. 5.**
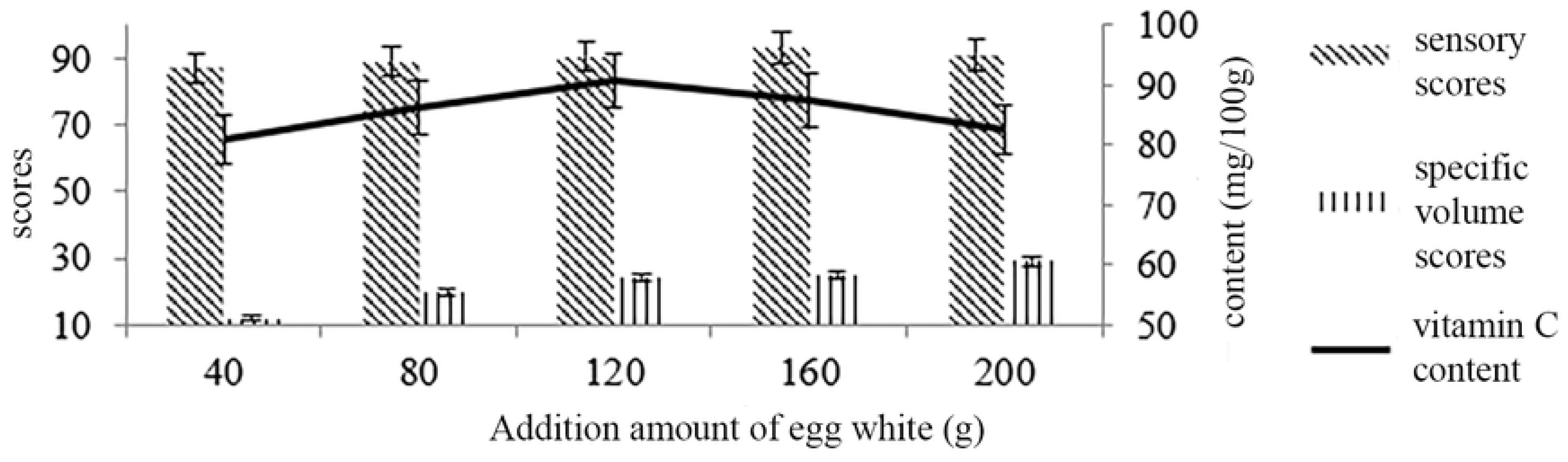
Effect of the amount of egg white on cake quality.

**S4 Table. Experimental results of response surface analysis.** The table shows the analysis results of the single-factor test.

**S5 Table.Variance analysis of regression equation.** In this table, “* * *” means extremely significant, “* *” means very significant, “and” * “means significant.”

**S6 Fig. 6.**
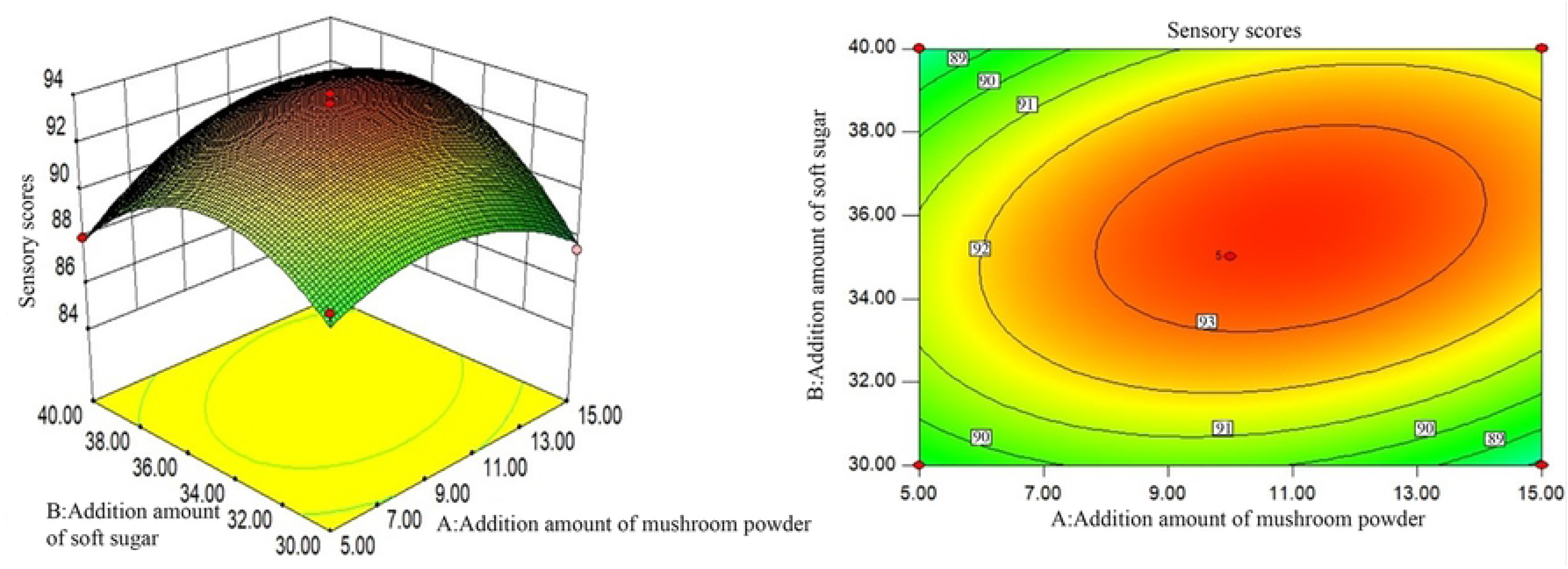
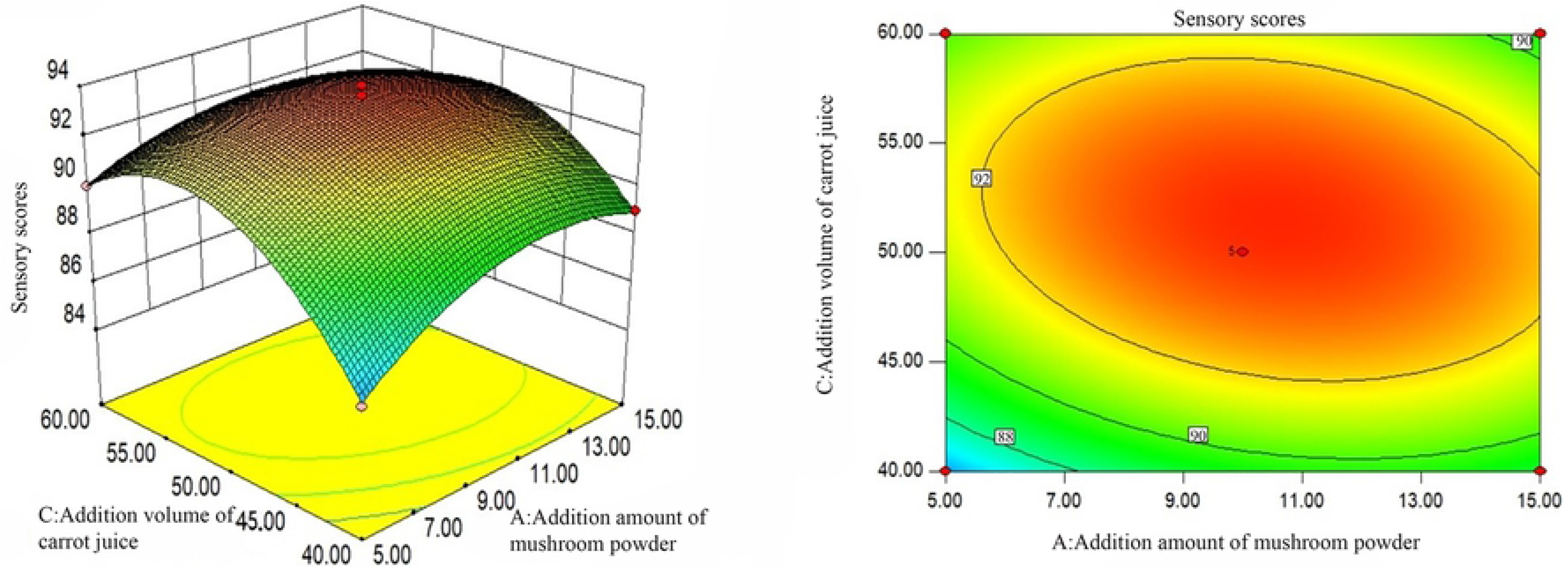
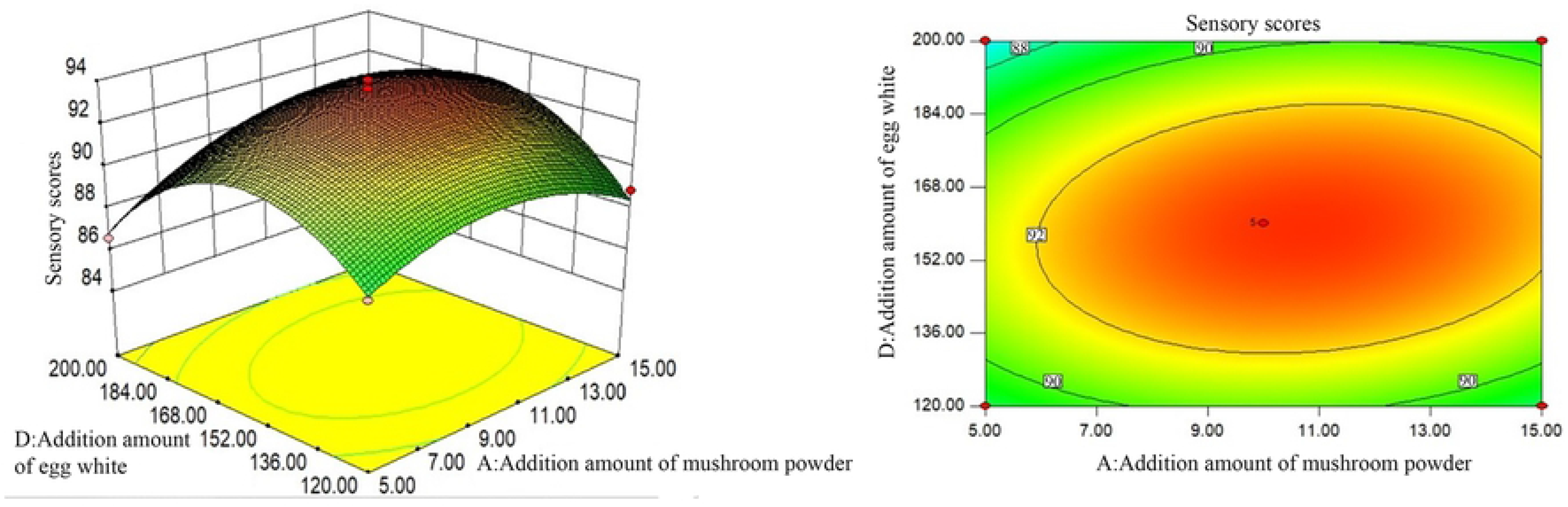
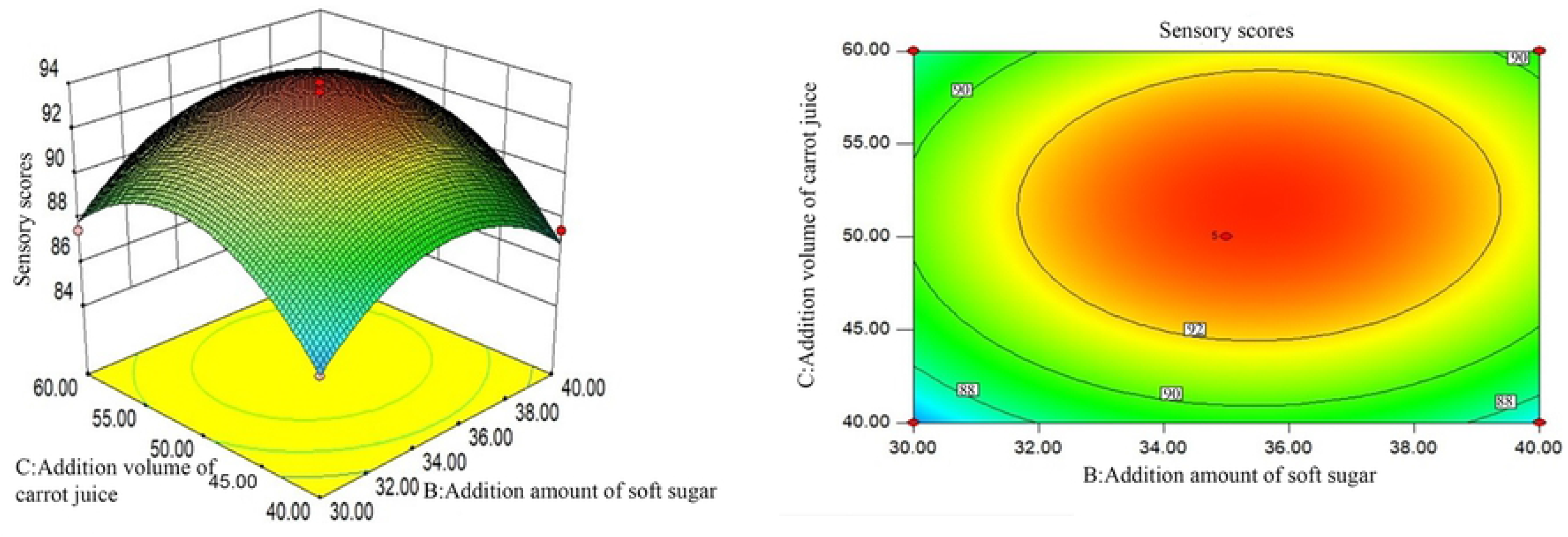
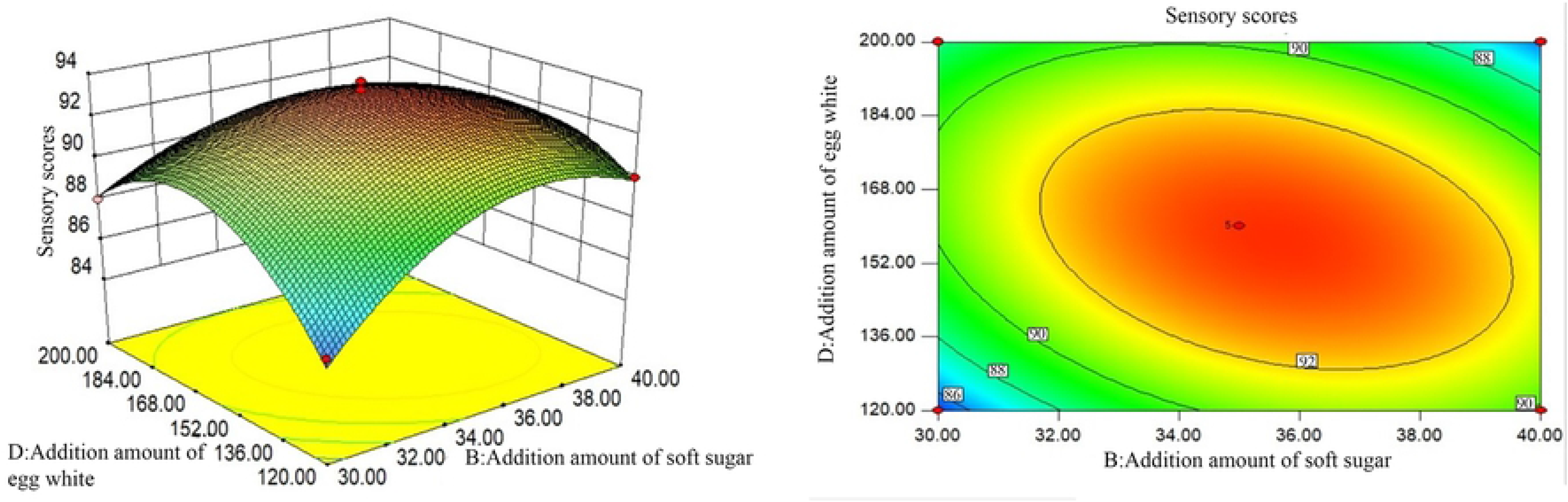
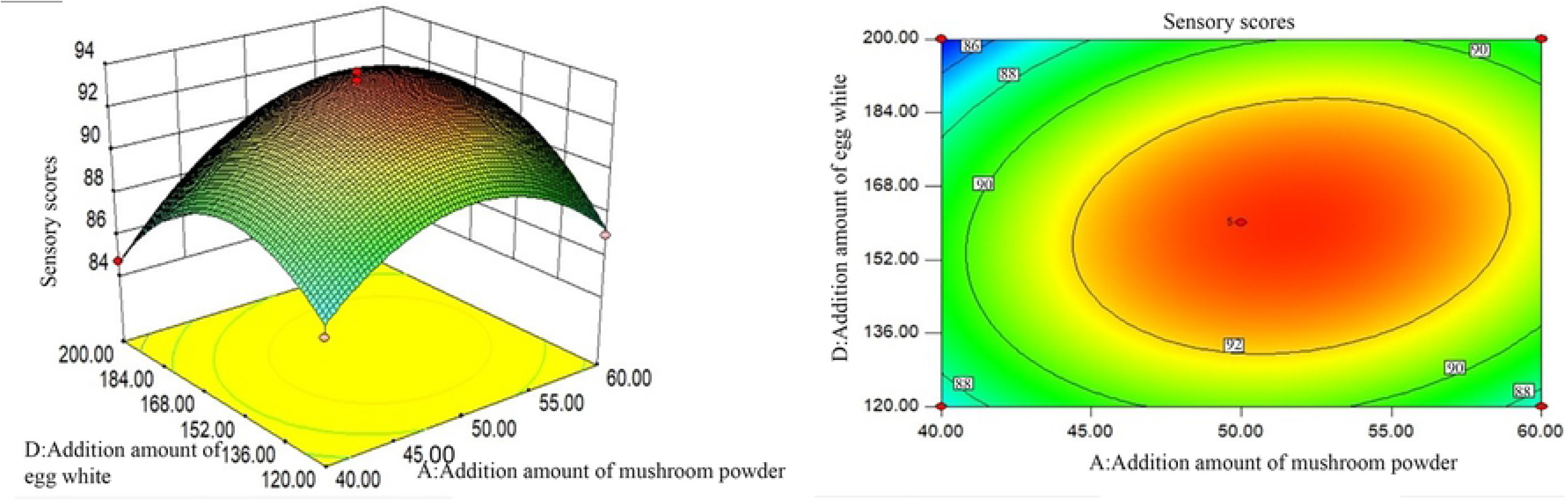
Effects of various factors on sensory scores of cake. This set of figures include response surface diagrams and contour line plots. (a) shows the effect of mushroom powder addition amount and soft sugar addition amount on sensory scores of cake. (b) shows the effect of mushroom powder addition amount and carrot juice addition volume on sensory scores of cake. (c) shows the effect of mushroom powder addition amount and egg white addition amount on sensory scores of cake. (d) shows the effect of carrot juice addition volume and soft sugar addition amount on sensory scores of cake. (e) shows the effect of soft sugar addition amount and egg white addition amount on sensory scores of cake. (f) shows the effect of carrot juice addition volume and egg white addition amount on sensory scores of cake.

**S7 Fig. 7.**
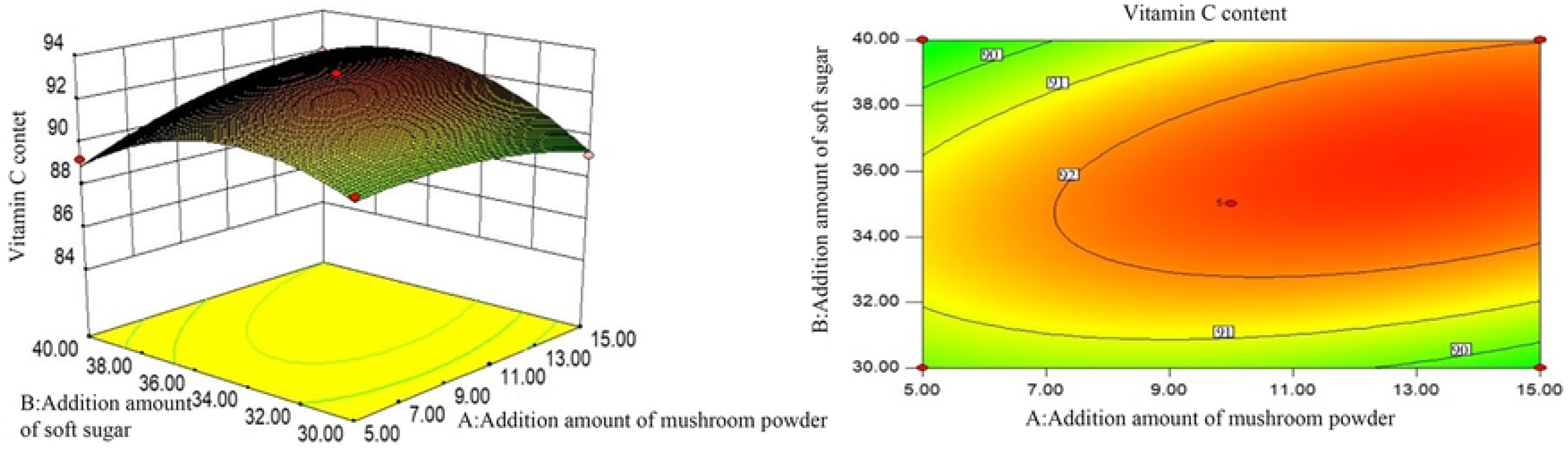
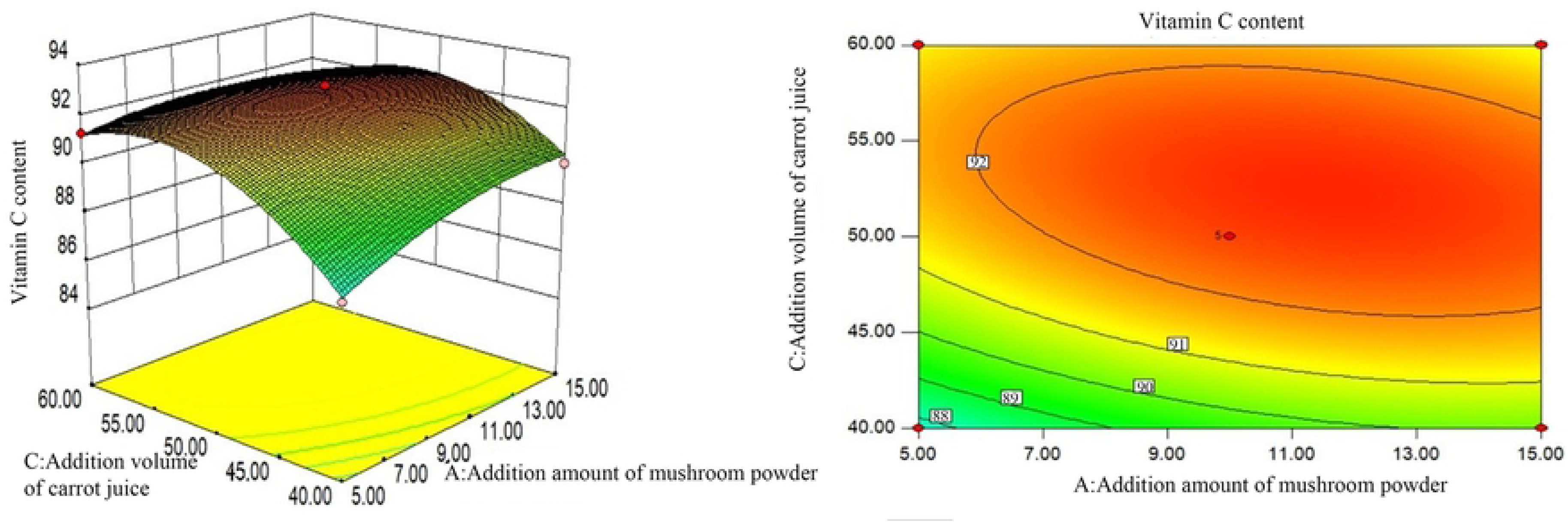
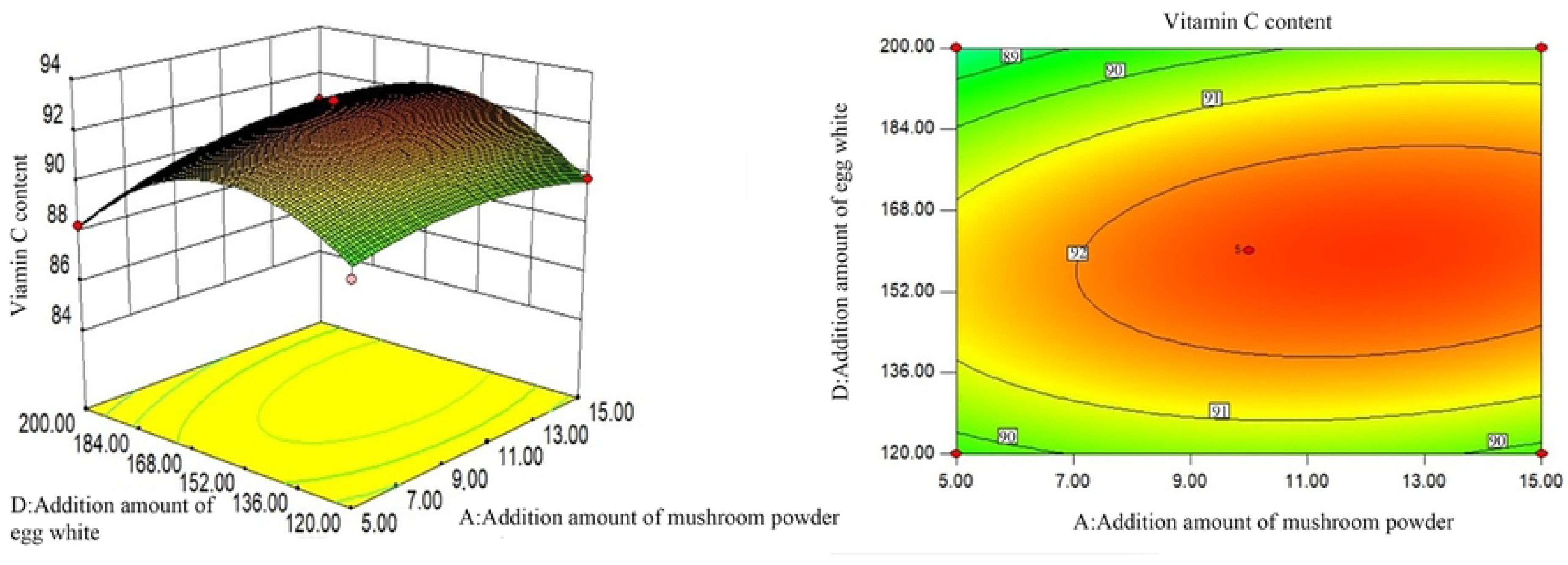
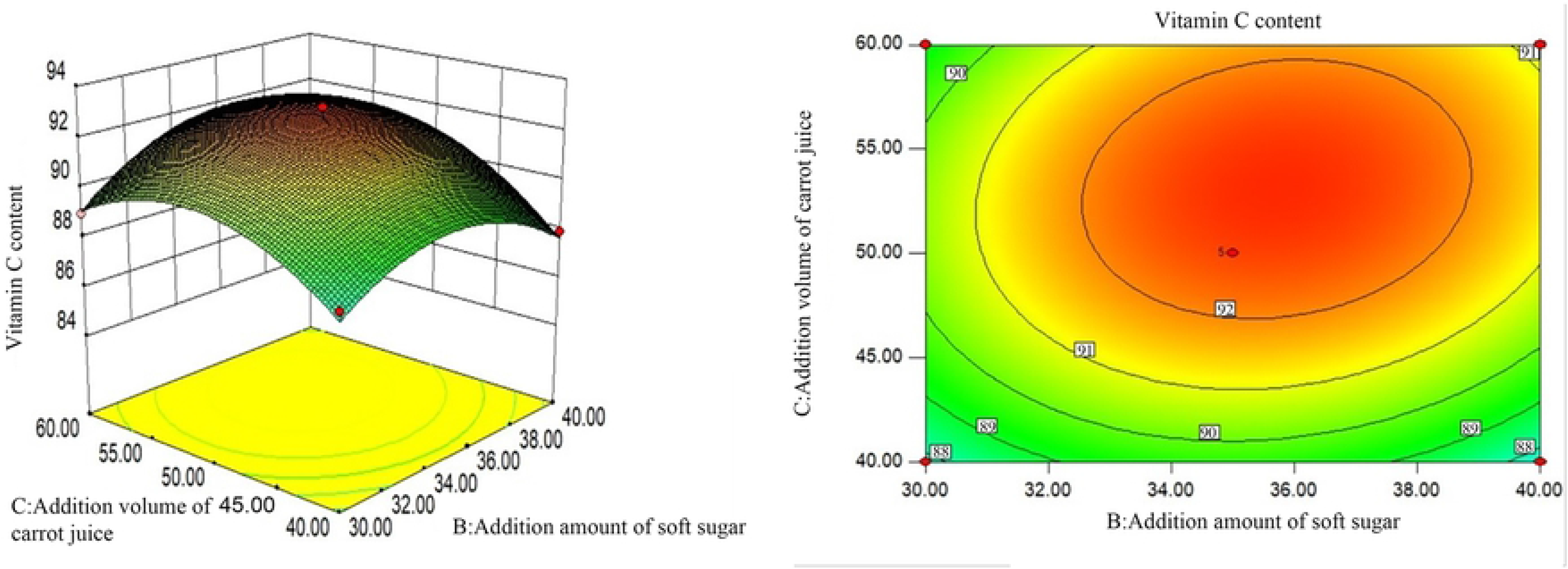
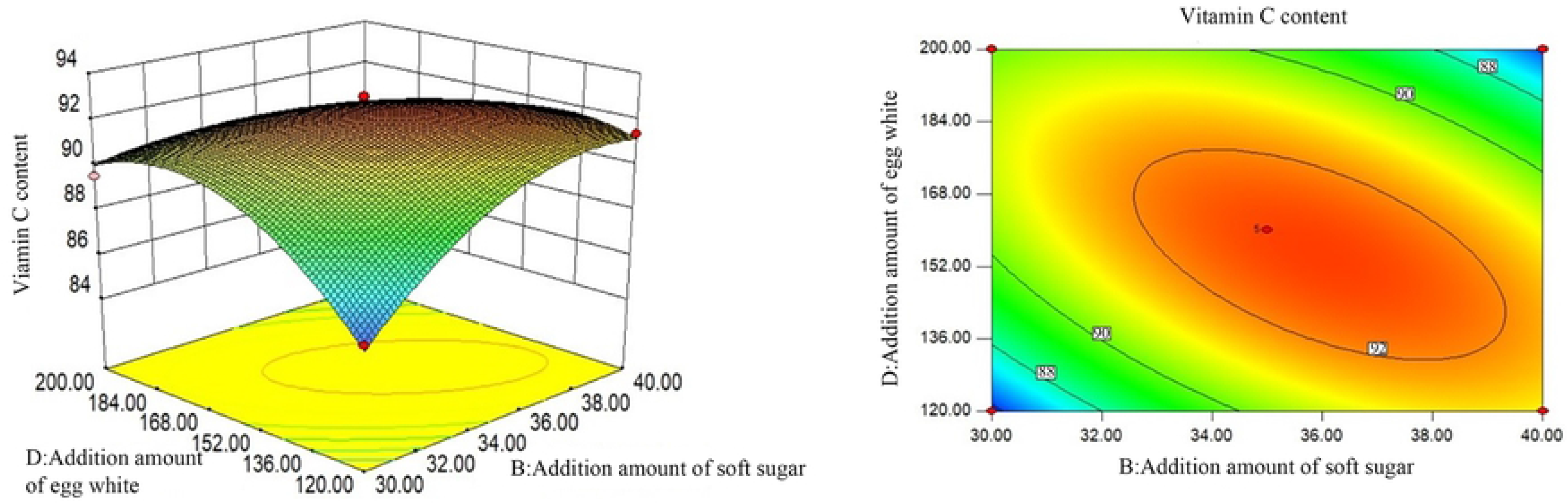
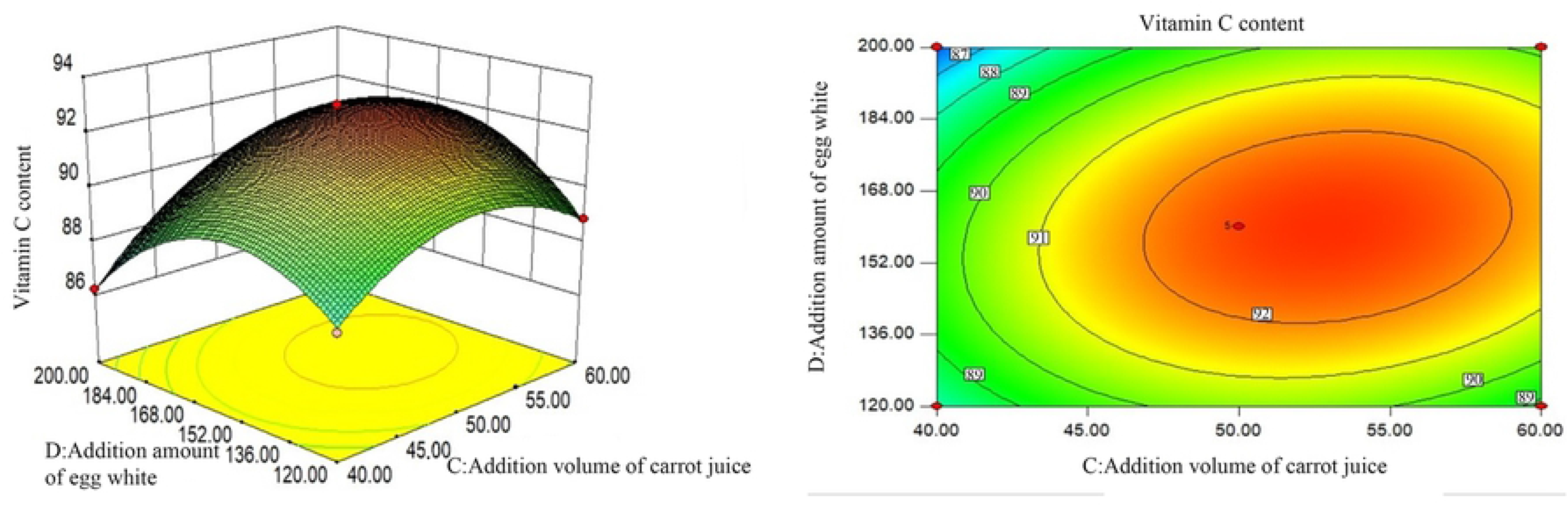
Effects of various factors on vitamin C content in cake. This set of figures include response surface diagrams and contour line plots, (a) shows the interactive effect of mushroom powder addition amount and soft sugar addition amount on vitamin C content in cake. (b) shows the interactive effect of mushroom powder addition amount and carrot juice addition volume on vitamin C in cake. (c) shows the interactive effect of mushroom powder addition amount and egg white addition amount on vitamin C content in cake. (d) shows the interactive effect of carrots juice addition volume and soft sugar addition amount on vitamin C content in cake. (e) shows the interactive effect of soft sugar addition amount and egg white addition amount on vitamin C content in cake. (f) shows the interactive effect of carrot juice addition volume and egg white addition amount on vitamin C content in cake.

